# Oxytocin neurons in the anterior and posterior paraventricular nucleus have distinct behavioral functions and electrophysiological profiles

**DOI:** 10.1101/2025.08.03.668226

**Authors:** Audrey N. Chrisman, Chiho Sugimoto, Hanna Butler-Struben, Vanessa A. Minie, Andrew L. Eagle, Daniela Anderson, Natalia Duque-Wilckens, Ashley Ramos, Yasmine I. Lewis, Sinéad C. Archdeacon, A. J. Robison, Brian C. Trainor

## Abstract

Oxytocin is a neuropeptide that can promote or inhibit affiliative social behaviors. Recent evidence suggests that these diverse effects are mediated by distinct oxytocin receptor-expressing neurons. An outstanding question is whether these behavioral effects are also driven by distinct or overlapping populations of oxytocin-producing neurons. The paraventricular nucleus (PVN) of the hypothalamus is a major source of oxytocin and sends projections to the mesolimbic dopamine system and extended amygdala. Previous work found that social defeat increased oxytocin neuron activity in the anterior PVN (aPVN) but not posterior PVN (pPVN). We reduced oxytocin synthesis with antisense morpholino oligonucleotides in either anterior or posterior PVN in California mice (*Peromyscus californicus*), a strong model system for studying effects of social stress on brain function and behavior. Antisense morpholinos in aPVN had no effect on behavior in unstressed females but increased social approach and reduced social vigilance in females exposed to social defeat stress. In pPVN, antisense morpholinos reduced social approach in unstressed male and female California mice. We then used *Oxt*^Cre^ mice to compare electrophysiological profiles of oxytocin in aPVN and pPVN with a population of oxytocin neurons in the bed nucleus of the stria terminalis (BNST). Oxytocin neurons in aPVN and BNST had higher post-synaptic events and responded more strongly to current injections than oxytocin neurons in pPVN, though they had similar excitatory and inhibitory input balance at the observed resting membrane potential. These findings shed light onto functional and physiological heterogeneity of PVN oxytocin neurons. Our results suggest that context dependent effects of oxytocin are mediated by different populations of oxytocin neurons.

## Introduction

The neuropeptide oxytocin is widely known to promote social behaviors in affiliative social contexts (Carter et al., 2020; Froemke and Young, 2021; Young and Wang, 2004). These effects have generated interest in potential therapeutic uses for oxytocin and similar neuropeptides (Neumann & Slattery, 2016; Bales et al., 2007). A challenge for implementing this approach is that data in humans (Eckstein et al. 2015; Quintana et al. 2021) and non-human animals (Beery, 2015; Moaddab and Dabrowska, 2017) show that oxytocin can increase anxiety-related behaviors. The social salience hypothesis provides a potential explanation for these divergent findings by positing that oxytocin enhances the salience of social cues across contexts (Bartz et al. 2011; Shamay-Tsorry & Abu-Akel, 2016). In affiliative contexts, oxytocin facilitates social approach responses (Preckel et al., 2014; Smith and Wang, 2014), while in agonistic contexts, oxytocin can drive avoidance and anxiety-like behavior (Declerck et al., 2010; Sofer et al., 2024). There is growing evidence that different populations of oxytocin receptors in distinct neural circuits have different effects on social behavior (Steinman et al. 2019; Guzman et al. 2013; Osakada et al. 2024). For example, oxytocin receptor antagonist infused into the nucleus accumbens (NAc) reduced social approach in non-stressed California mice (Williams et al., 2020) and pair bond formation in prairie voles (Liu and Wang, 2003). In contrast, oxytocin receptor agonist infused into the anteromedial bed nucleus of the stria terminalis (BNST) reduced social approach and increased vigilance non-stressed California mice (Duque-Wilckens et al., 2020; Luo et al., 2022). It remains unclear whether these distinct populations of oxytocin receptor-expressing neurons receive input from different populations of oxytocin-expressing neurons. For example, oxytocin neurons within the BNST are activated when animals are expressing social avoidance responses (Nasanbuyan et al., 2018; Steinman et al., 2016), and oxytocin knockdown within the BNST reverses stress-induced social avoidance (Duque-Wilckens et al., 2020). However, it is unknown if BNST oxytocin neurons are unique in promoting social avoidance responses, given that the paraventricular nucleus (PVN) of the hypothalamus sends oxytocin fibers to the BNST (Eliava, et al. 2016).

The PVN contains more oxytocin neurons than the BNST, and these neurons project to numerous brain regions that impact social behavior, including the NAc (Knobloch et al., 2012; Son et al., 2022). Oxytocin neurons are traditionally classified as magnocellular or parvocellular (Swanson and Sawchenko, 1980), but recent findings suggest that this binary categorization may be too simplistic (Althammer and Grinevich, 2018). For example, an analysis of morpho-electric properties of PVN oxytocin neurons in C57Bl6/J mice (Chen et al., 2022) revealed some magnocellular neurons in the posterior PVN (pPVN) are more similar to pPVN parvocellular neurons than to magnocellular neurons in anterior PVN (aPVN). Some evidence suggests that a functional anterior-posterior organization of oxytocin neurons within the PVN may contribute to regulation of distinct types of social behavior. In juvenile C57BL6/J mice, oxytocin neurons that promote social learning are clustered in the pPVN (Dölen et al., 2013; Lewis et al., 2020). Conversely, in female California mice (*Peromyscus* californicus) examined two weeks after defeat stress, had increased oxytocin/c-fos colocalizations in the aPVN but not pPVN in mice that exhibited social avoidance and vigilance (Steinman et al., 2016). These findings support the idea that distinct populations of oxytocin neurons within the PVN may promote social approach versus social vigilance and avoidance responses.

To test this hypothesis, we used a morpholino antisense knockdown strategy to selectively inhibit oxytocin production in the aPVN or pPVN and examined their effects on social behavior. We performed behavioral studies on California mice, a territorial and monogamous rodent in which males and females defend joint territories (Ribble et al., 1990). This behavioral organization has made this species ideal for studying the impact of social stress in both sexes (Lake and Trainor, 2024). Social defeat induces lasting (2+ weeks) reductions in social approach in female but not male California mice (Trainor et al., 2013; Wright et al., 2023), and defeat increases the reactivity of aPVN oxytocin neurons in females but not males in a social interaction test at the same 2-week timepoint (Steinman et al., 2016). Building on these findings, we tested whether oxytocin knockdown affected behavior in control or stressed females. We previously observed no sex differences in how pPVN oxytocin neurons responded during social interactions tests, so we examined how oxytocin knockdown affected behavior in stress naïve males and females. To couple behavioral output to functional differences, we complemented these studies with electrophysiological analyses of oxytocin neurons in *Oxt*^Cre^ C57Bl/6J mice (Carcea et al., 2021; Marlin et al., 2015). Together, these analyses suggest that the electrophysiological and functional properties of aPVN oxytocin neurons more closely resemble BNST oxytocin neurons than pPVN neurons.

## Methods

### Subjects

For behavioral experiments, California mice (*Peromyscus californicus)* were bred and maintained in a colony at UC Davis on a 16L:8D light-dark cycle with *ad libitum* access to food (Harlan Teklad) and water. The mice were housed in clear propylene cages with wire bar tops in groups of same-sex groups of 2-4. The cages were bedded with Sani-Chip (Harlan Laboratories, Indianapolis, IN, USA), Enviro-Dri (Eco-bedding, Fibercore, Cleveland, OH, USA), and Nestlets (Ancare, Bellmore, NY, USA). All animals were at least 90 days old at the start of the study. All behavioral assessments were conducted during the dark phase under dim red light (3 lux). For electrophysiology experiments, Oxt-IRES Cre mice (Oxt^Cre^, Strain #024234) were purchased from Jackson Labs and crossed with Cre-inducible Rosa^eGFP-L10a^ mice (Gift of Dr. Gina Leinninger at Michigan State University). All procedures followed NIH guidelines and were approved by UC Davis and Michigan State University Institutional Animal Care and Use Committees.

### Electrophysiology

*Ex vivo* whole cell slice electrophysiology was conducted in *Oxt*^Cre^/Rosa^L10eGFP^ mice aged 7-10 weeks. All solutions were bubbled with 95% O_2_ -5% CO_2_ throughout the procedure. Mice were anesthetized with isofluorane and transcardially perfused with ice cold sucrose artificial cerebrospinal fluid (aCSF) (234 mM sucrose, 26 mM NaHCO_3_, 11 mM D-glucose, 10 mM MgSO_4_, 2.5 mM KCl, 1.25 mM NaH_2_PO_4_, 0.5 mM CaCl_2_). Brains were rapidly removed, blocked, and sectioned into 250 μM coronal sections containing the brain region(s) of interest using a vibratome (Leica VT1200S R). Brain slices were transferred to an incubation chamber containing saline aCSF (126 mM NaCl, 26 mM NaHCO_3_, 10 mM glucose, 2.5 KCl, 2 mM MgCl_2_, 1.25 mM NaH_2_PO_4_) held at 37° C for 30 min, then at room temperature until used for recordings. Recordings were made from slices held in a submersion chamber perfused with saline aCSF (2 mL/min) held at 32 °C with an inline heater (Warner Instruments). Borosilicate glass electrodes (3–6MΩ) were filled with a potassium gluconate internal solution (115 mM potassium gluconate, 20 mM KCl, 10 mM phosphocreatine-di(Tris), 2 mM Mg-ATP, 1.5 mM MgCl_2_, 0.5 mM Na 3 -GTP; pH 7.2; 285–295 mOsm). GFP-positive cells were visualized using an Olympus BX51WI microscope using DIC infrared and epifluorescent illumination. Whole-cell patch-clamp recordings were made from cells using a Multiclamp 700B amplifier and Digidata 1440A digitizer (Molecular Devices) and whole-cell junction potential was not corrected. Recordings were sampled (10 kHz), filtered (10 kHz), and digitally stored. Membrane capacitance, and membrane resistance were automatically calculated by pClamp 10 software (Molecular Devices). Resting membrane potential was measured by the Multiclamp without injecting current (I=0). Spontaneous PSC event frequency and amplitude were determined from gap free voltage clamp recording and analyzed using MiniAnalysis software (Synaptosoft, Inc.) and Clampfit Minis Search software (Molecular Devices). Excitability of neurons was measured by increasing depolarizing steps (0 to 100 pA, Δ25 pA steps, 500 ms) with 30 s step intervals. Presence of a transient outward rectification, which is found in magnocelluar oxytocin neurons but not parvocellular oxytocin neurons was used to differentiate the neuron types. To find this transient outward rectification, a minimum current to reach ∼-100 mV (200 ms) was applied followed by increasing depolarizing steps (200 ms, Δ25 pA) with a 2 s sweep interval.

### Social Defeat Stress

Female California mice were randomly assigned to social defeat stress or a control handling condition (Greenberg et al., 2014; Trainor et al., 2013). In the social defeat stress group, mice were placed in the home cage of a highly aggressive same-sex counterpart and were either exposed to 7 bites from the resident or remained in the cage for 7 minutes, whichever occurred first. Mice assigned to the control group were placed into an empty cage for 7 minutes. Social defeat sessions and control handling sessions occurred for three consecutive days. After each episode, all mice were returned to the home cage.

### Morpholino inhibition of oxytocin synthesis

A morpholino stock solution (Genetools, LLC) was created using artificial cerebrospinal fluid to create a 150 pM solution. The antisense sequence targeting oxytocin mRNA, previously validated in the BNST (Duque-Wilckens et al., 2020), was 5′-TTG GTG TTC TGA GTC CTC GAT CC-3′. The missense control sequence was 5′-TTC GTC TTC TGA CTC CTC CAT GC-3′. California mice were randomly assigned to receive a 0.2 μL infusion of either antisense or missense solution. Injections were made either into the aPVN (Fig. 1, coordinates: anterior-posterior: -0.8, medial-lateral: ±0.3, dorsal-ventral: −6.1) or pPVN (Fig. 1, coordinates: anterior-posterior: -1.2, medial-lateral: ±0.7, dorsal-ventral: −5.1) using a sterile glass pipette. Carprofen (5 mg/kg) was administered subcutaneously immediately following surgery and daily for the next 3 days. Six days after surgery, behavior was assessed in the social interaction test.

**Figure 1.**
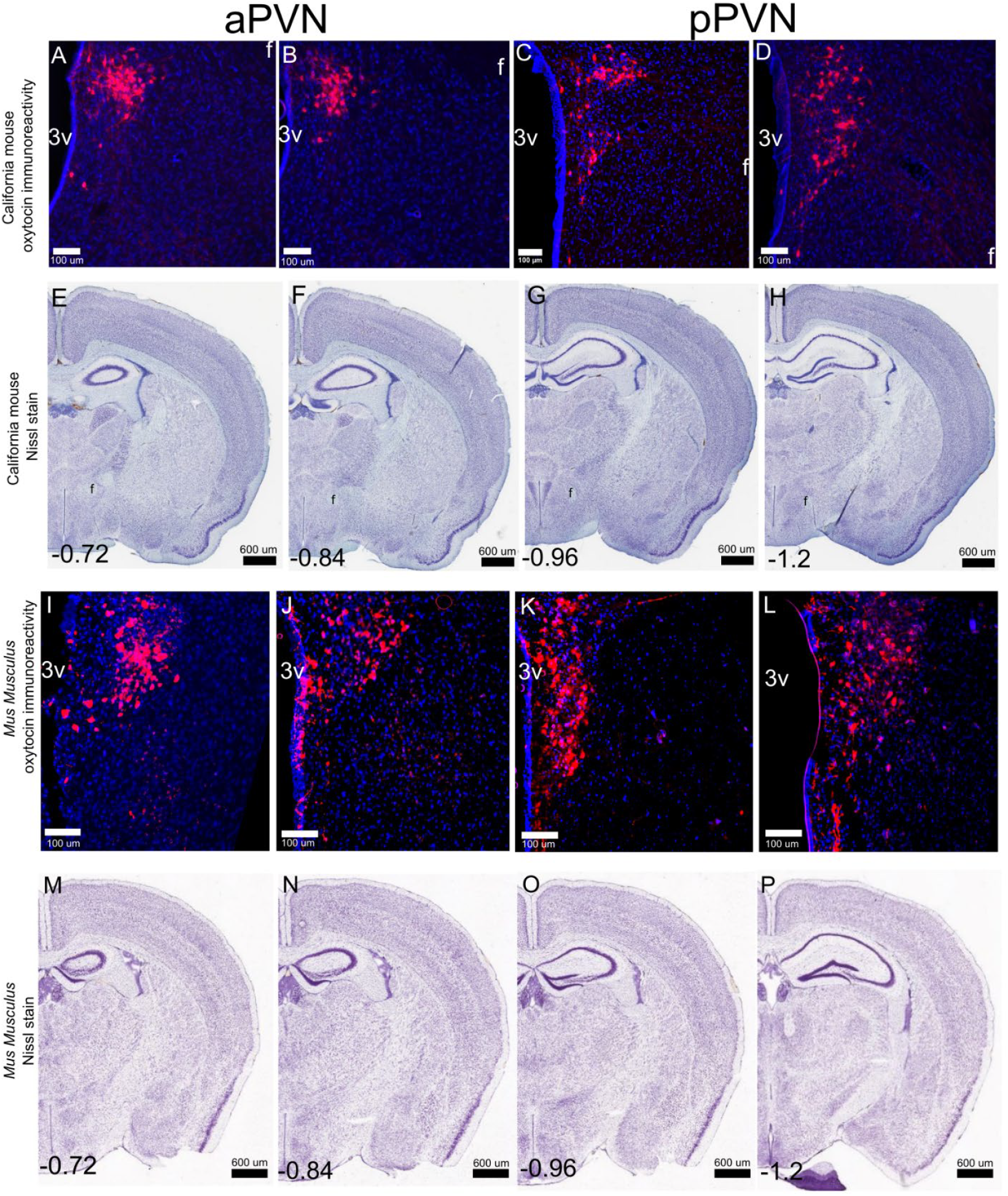
Distribution of oxytocin neurons (red) across the anterior-posterior extent of the paraventricular nucleus in California mice (A-D) and Mus Musculus (I-L) with corresponding Nissl images from the Brainmaps California mouse brain atlas (J-L) (brainmaps.org) and Allen mouse brain atlas (M-P).

### Social interaction test

The social interaction test consisted of three consecutive three-minute phases: open field, habituation, and interaction. In the open field phase, the mouse roamed freely in the 89 cm x 63 cm x 60 cm arena. In the acclimation phase, a 22cm x 15 cm x 15 cm wire cage was introduced into the back of the arena to familiarize the mouse with the new structure before introducing the target mouse. In the interaction phase, a novel conspecific target mouse of the same sex was placed into the wire cage inside the arena, allowing the mouse to interact with the caged target mouse. An overhead camera and ANY-maze software (Stoelting) recorded and tracked the animal’s locomotion, time spent in the interaction zone (within one body length of the wire cage), and time spent in the arena’s center. Vigilance behavior in videos of the interaction phase was manually scored by the time spent outside the interaction zone and with the head turned to the target cage. The time spent engaging in vigilance out of the three-minute trial was quantified using BORIS (Friard & Gamba, 2016) and averaged.

### Tissue collection for histology

For California mice, one day following the social interaction test, animals were euthanized using isoflurane, then perfused with 4% paraformaldehyde (PFA). The brains were collected, stored in 4% PFA overnight, then switched to 30% sucrose solution for a minimum of 48 hours. Following sucrose treatment, the brains were flash frozen with dry ice then immediately stored in -80° C. Frozen brains were sliced coronally with a cryostat (Leica) at -20° C at 40 mm and stored in cryoprotectant. For *Oxt*^Cre^ validation studies, mice were deeply anesthetized and transcardically perfused with ice cold PBS following by 10% formalin. Brains were fixed overnight with 10% formalin then stored in 30% sucrose until sectioned to 35 mm using a freezing microtome (Leica SM2010R).

### Immunohistochemistry

For California mice coronal slices at 120 µm intervals containing the PVN were selected. After removal from storage in cryoprotectant, the slices were washed twice for 5 minutes in phosphate-buffered saline (PBS) before being blocked with 10% normal goat serum. Then, the tissue incubated with mouse anti-oxytocin primary antibody (1:2000; Millipore Sigma PS38) in 2% normal goat serum for 24 hours at 4° C. This primary antibody has been validated for specificity in California mice (Steinman et al., 2015). After primary antibody incubation, the slices were washed three times for 5 minutes in PBS to remove all unbound antibodies before incubating with secondary antibody goat anti-mouse Alexa-Fluor 555 (1:500, ThermoFisher) in 2% normal goat serum for 2 hours at room temperature. After another 5-minute wash in PBS, sections were mounted onto glass slides and coverslipped with mounting medium (Vectashield).

For *Oxt*^Cre^ mice, brain slices were washed with 0.3% Tween20 in PBS three times for 10 minutes, then blocked with 10% normal donkey serum before incubated in primary antibodies 1:4000 goat anti-GFP (Abcam, ab5450) and 1:1000 mouse anti-neurophysin1 (Millipore Sigma PS38) in 10% NDS and 0.6% TritionX100 in PBS overnight. After primary antibody incubation, the slices were washed three times for 5 minutes in PBS to remove all unbound antibodies before incubating with secondary antibody 1:500 secondaries donkey anti-goat AF488 and anti-mouse Cy3 in 1% normal donkey serum and 0.3% Tween20 in PBS. Sections were mounted onto glass slides and coverslipped with DPX mounting media (Sigma). Fluorescent images were taken using a Nikon Eclipse Ni-U Upright Fluorescent Microscope.

### Quantification of oxytocin knockdown

Photomicrographs of the tissue were captured using a Keyence BZX-800. We used ImageJ (NIH, Bethesda, MD) to analyze percent staining for each image. An observer who was unaware of treatment group assignment counted cell bodies with positive oxytocin staining. Anterior PVN was defined as -0.8 to -0.99 Bregma and pPVN was defined as -1.0 to -1.2 Bregma.

## Results

### Oxytocin neurons in anterior and posterior PVN

Coronal sections across the anterior-posterior extent of the PVN were used to characterize the distribution of oxytocin neurons in anterior versus posterior PVN. In aPVN (**Fig. 1A-C, 1F-H**), oxytocin positive neurons are located below the dorsal extent of the 3^rd^ ventricle (3v), and the fornix is positioned lateral and dorsal to these cells. In the pPVN, oxytocin neurons are positioned adjacent to the dorsal border of the 3v and the fornix is lateral and ventral (**Fig 1D-E, 1I-J**).

### Knockdown of oxytocin in the anterior PVN

Previous analyses of oxytocin/c-fos colocalizations suggested distinct patterns of activation in anterior vs posterior PVN (Steinman et al. 2016), so we used morpholinos to reduce oxytocin production in aPVN (**Fig. 2A**). Antisense injected into the aPVN reduced the number of oxytocin-immunoreactive neurons in aPVN (t_31_=2.12, p<0.05) but not pPVN (**Fig. 2E**). In female California mice, the effects of oxytocin knockdown on social approach were dependent on stress exposure (**Fig. 2F**, stress*treatment F_1,47_=3.82, p =0.05). In missense treated females, social approach was significantly lower following stress exposure (**Fig.2F**, p<0.001,Cohen’s d=1.19). Stressed females treated with antisense showed significantly more social approach than stressed females treated with missense (**Fig. 2F**, p<0.05, d=0.76). Similar patterns were observed for social vigilance. Stressed females treated with missense had increased vigilance compared to control females treated with missense (**Fig. 2F**, Mann-Whitney p<0.05. d=0.71). Stressed females treated with antisense had significantly lower levels of social vigilance compared to stressed females treated with missense (**Fig. 2F**, Mann-Whitney p<0.05, d=0.76). There were no differences in time spent interacting with an empty cage (**Fig. 2E**), time spent in the center of the arena during the open field phase (**Fig. 2F**), or general locomotion (**Fig. 2G**). Together, these data suggest that oxytocin neurons in the aPVN have similar functions to oxytocin neurons in the BNST.

**Figure 2.**
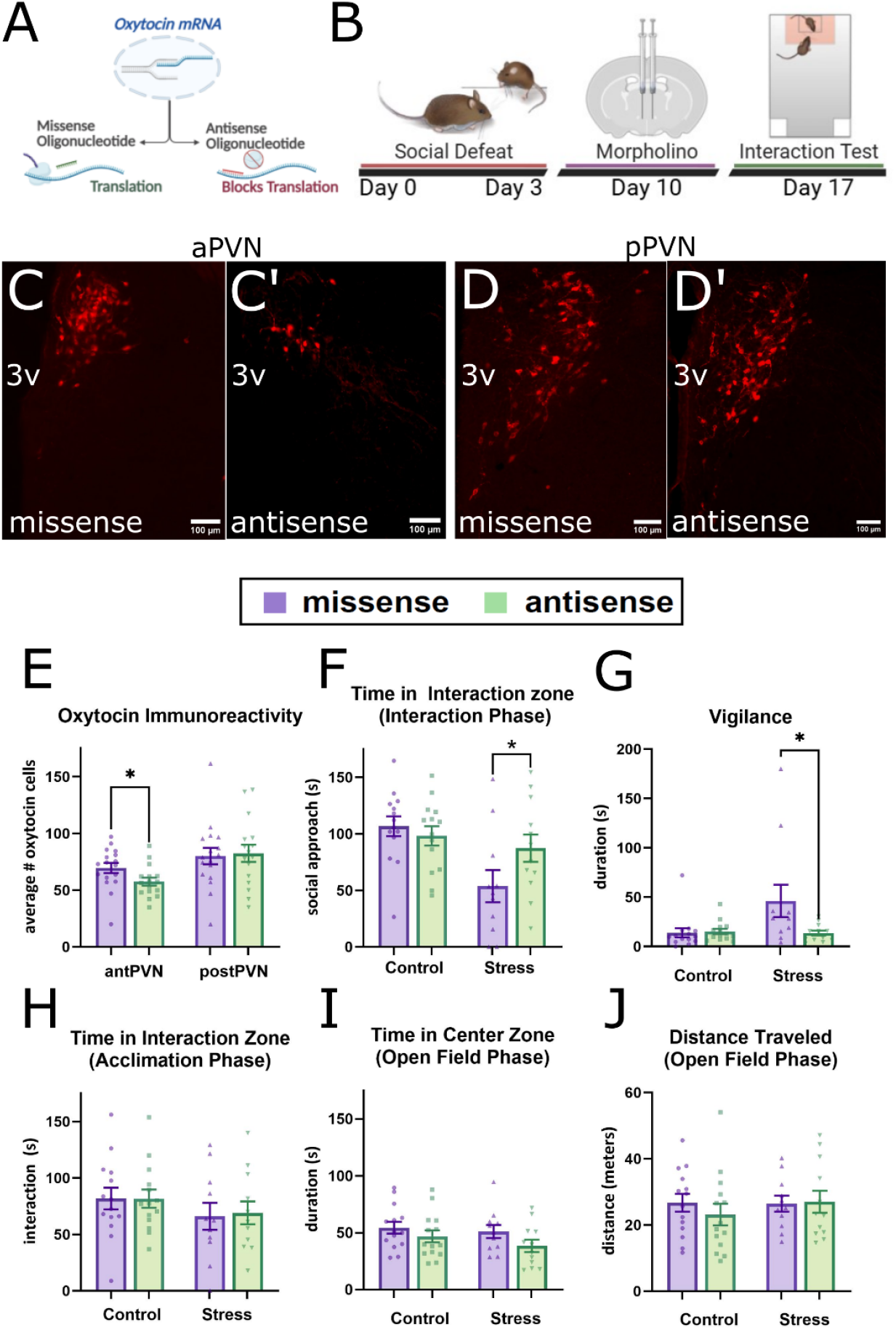
Effects of targeted knockdown of oxytocin via morpholinos in the anterior PVN of socially defeated or control handled female California mice. (A) Morpholinos block translation of oxytocin mRNA. (B) Experimental timeline. (C) Representative photomicrographs of oxytocin immunostaining in the anterior PVN of (C) missense-treated and (C’) antisense-treated mice. (D) Representative photomicrographs of oxytocin immunostaining in the posterior PVN of (D) missense-treated and (D’) antisense-treated mice. (E) Oxytocin immunoreactivity in the anterior and posterior PVN of missense and antisense-treated mice. * p<0.05 antisense anterior vs missense anterior. (F) Effects of morpholino treatment on time spent in the interaction zone with a novel conspecific in stressed and control females, (G) vigilance behavior, (H) time spent in the interaction zone with a novel empty cage, (I) time spent in the center zone during an open field test, and (J) distance traveled during an open field test. * p < 0.05 vs stress missense. N=11-15 per group.

### Knockdown of oxytocin in the posterior PVN

In pPVN, oxytocin morpholino treatment had different behavioral effects (**Fig. 3A**). Antisense injected into the pPVN reduced the number of oxytocin-immunoreactive neurons in pPVN (t_31_=5.89, p<0.001) but not aPVN (**Fig. 2E**). In unstressed males and females, antisense morpholino treatment reduced social approach (**Fig. 3B**, main effect F_1,31_=7,54, p=0.01). There were no significant sex differences or sex*treatment interactions. Morpholino treatment did not affect social vigilance during the social interaction phase (**Fig. 3C**), interaction with the empty cage during acclimation (**Fig. 3D**), time spent in the center of the arena (**Fig. 3E**), or general locomotion (**Fig. 3F)** during an open field test.

**Figure 3.**
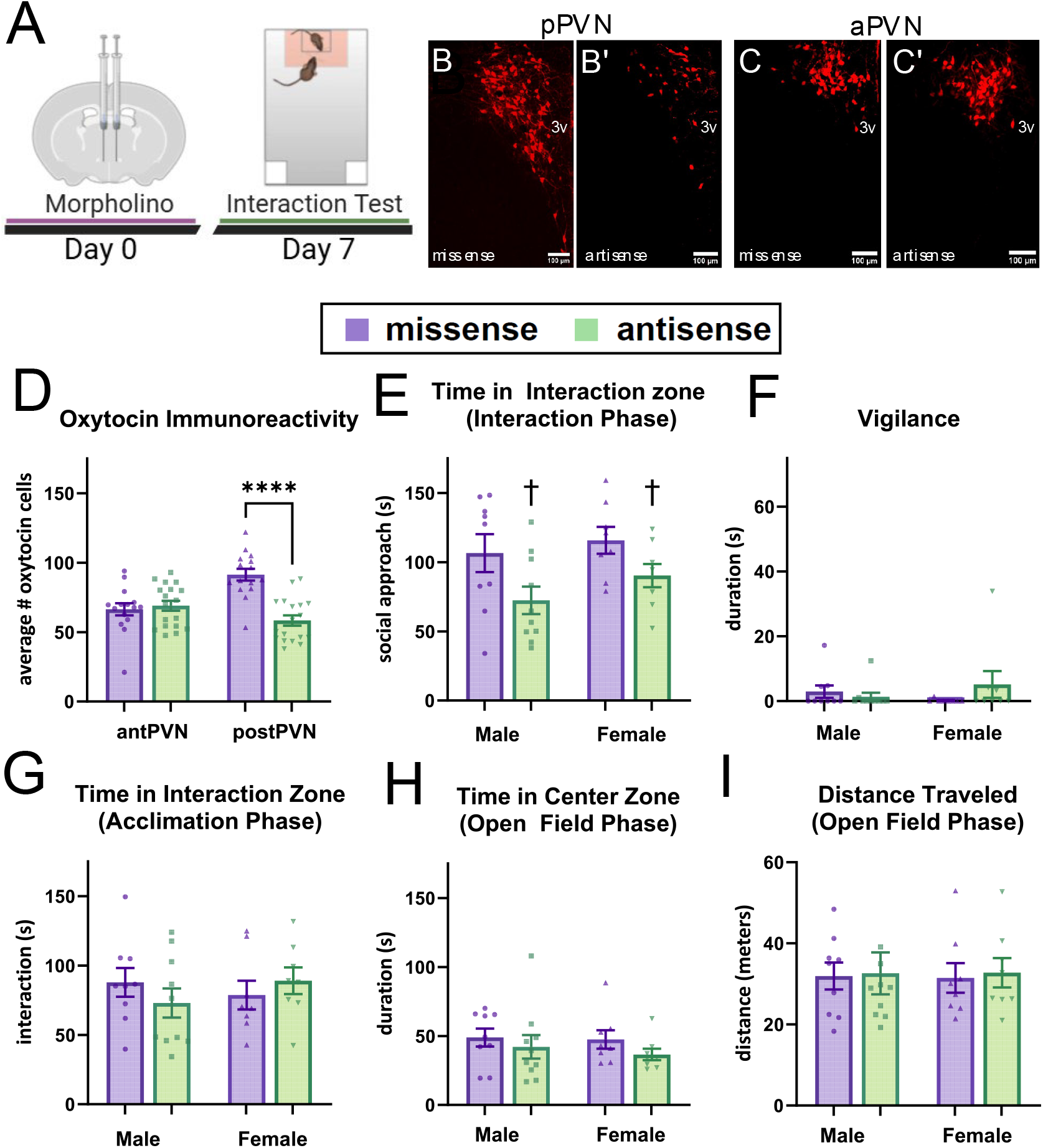
Effects of targeted knockdown of oxytocin via morpholinos in the posterior PVN of California mice. (A) Experimental timeline. (B) Representative photomicrographs of oxytocin immunostaining in the posterior PVN of missense-treated (B) and antisense-treated (B’) mice. (C) Representative photomicrographs of oxytocin immunostaining in the anterior PVN of missense-treated (C) and antisense-treated (C’) mice. (D) Oxytocin immunoreactivity in the anterior and posterior PVN of missense and antisense-treated mice. (E) Effects of morpholino treatment on time spent in the interaction zone with a novel conspecific in stressed and control females, (F) vigilance behavior, (G) time spent in the interaction zone with a novel empty cage, (E) time spent in the center zone during an open field test, and (F) distance traveled during an open field test. N=11-15 per group. † p < 0.01 vs main effect of antisense treatment. **** p<0.001 vs pPVN missense.

### Electrophysiological characterization of oxytocin neurons

To characterize intrinsic electrophysiology properties of oxytocin neurons across the PVN and BNST, we performed whole-cell patch clamp recordings from GFP-containing oxytocin and GFP-negative non-oxytocin neurons in the *Mus musculus* PVN, BNST, and cortex. Using immunohistochemistry, we found that GFP (**Fig. 4A**) was selectively expressed in in oxytocin positive neurons (**Fig. 4B**,**C**). Interestingly, oxytocin neurons in the pPVN had depolarization kinetics similar to parvocellular oxytocin neurons in Wistar rats (Eliava et al., 2016). Specifically, these cells lacked a transient outward rectification (**Fig. 4F**) that has been observed in magnocellular oxytocin neurons (Luther et al., 2002). We observed this distinctive feature in aPVN oxytocin neurons (**Fig. 4G**) as well as BNST oxytocin neurons (**Fig. 4H**). Cortical neurons (**Fig. 4I**) and non-oxytocin neurons did not exhibit transient outward rectification.

**Figure 4.**
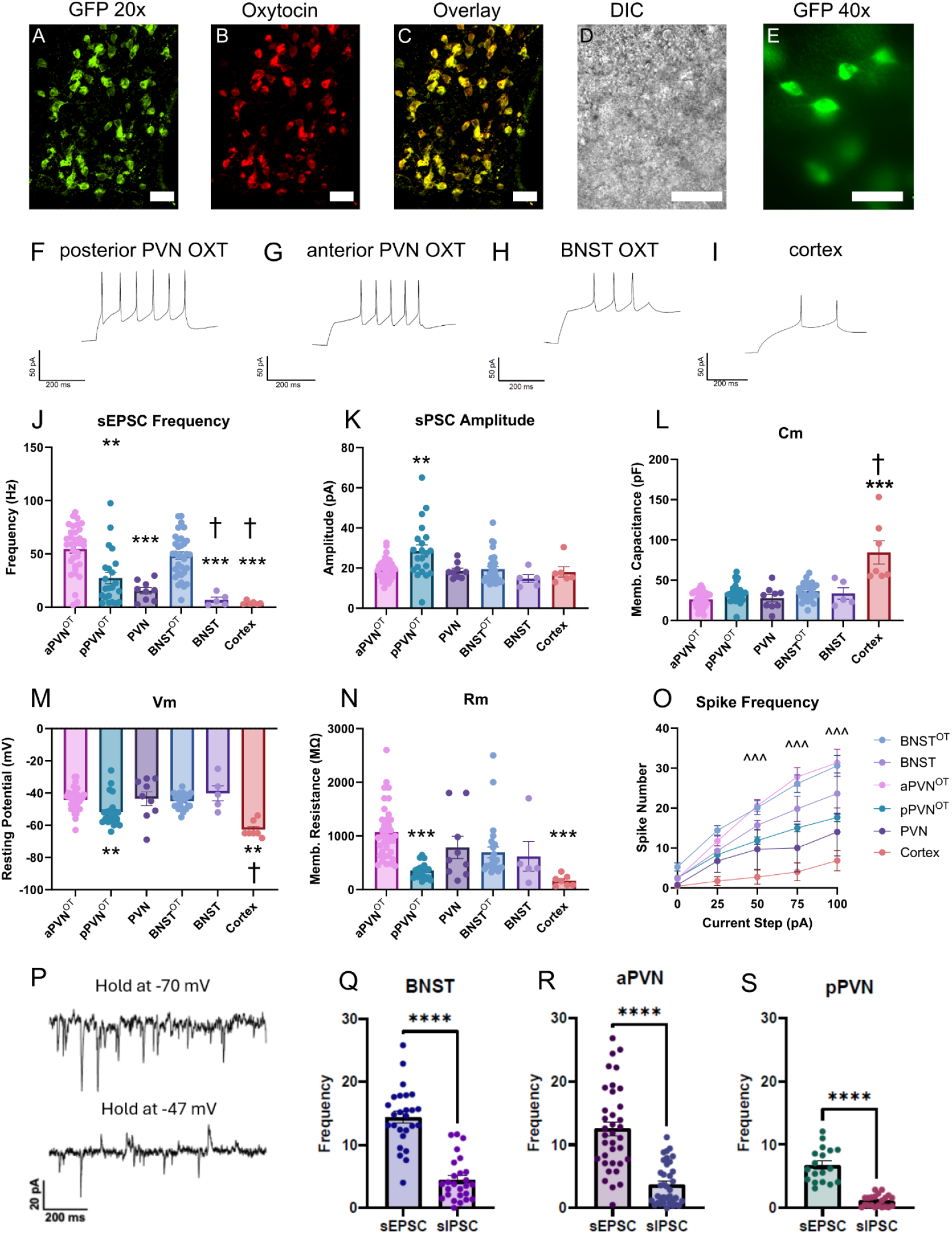
Electrophysiological characteristics of oxytocin and non-oxytocin neurons in the BNST and PVN of *Oxt*^Cre^/Rosa^L10eGFP^ mice. Representative images of: A) GFP (20x) B) Oxytocin immunostaining (20x) C) GFP-Oxytocin overlay (20x) D) Differential interference contrast (DIC) image (40x) E) GFP (40x). Representative current-clamp recordings from: F) posterior PVN oxytocin neurons (pPVN^OT^), G) anterior PVN oxytocin neurons (aPVN^OT^), H) BNST oxytocin neurons (BNST^OT^), and I) cortical neurons. Electrophysiological properties across groups are quantified for: J) sPSC frequency K) sPSC amplitude L) membrane capacitance M) resting membrane voltage N) membrane resistance O) spike frequency in response to 0-150 pA current injection. (P) Example current traces showing that oxytocin neurons show both upward and downward events when held at -47 mV. Q) Frequency of sEPSC and sIPSC in the oxytocin neurons of the (Q) BNST, (R) aPVN, and (S) pPVN. ** p<0.01 vs aPVN, *** p<0.01 vs aPVN^OT^, **** p<0.001 vs sIPSC † p<0.01 vs BNST^OT^ ^^^ p<0.001 aPVN^OT^ vs pPVN^OT^. For cell propagation analyses a total of 111 cells (64 male, 47 female) were recorded from 35 mice (19 male, 16 female). For sPSC analyses a total of 133 cells (77 male, 56 female) were recorded from 32 mice (16 male, 16 female). For spike frequency analyses a total of 92 cells (49 male, 43 female) were recorded from 28 mice (15 male, 13 female).

The frequency of spontaneous postsynaptic currents (sPSCs) differed across brain regions and cell types (**Fig. 4J**, F_5,103_=15, p<0.0001). Oxytocin neurons in the pPVN (pPVN^OT^) had significantly lower sPSC frequencies than oxytocin neurons in the aPVN (aPVN^OT^; planned comparison p<0.0001). Anterior PVN oxytocin neurons also had higher sPSC frequencies than non-oxytocin PVN neurons (p<0.001). These data suggest that oxytocin neurons in the aPVN have more robust synaptic input than pPVN oxytocin neurons.

Similarly, BNST oxytocin (BNST^OT^) neurons had higher sPSC frequencies compared to non-oxytocin BNST neurons (p<0.01). In this respect, BNST and aPVN oxytocin neurons appear to share strong synaptic input or regulation. Cortical neurons recorded on the same slices as PVN neurons had lower sPSC frequencies than aPVN oxytocin (p<0.01) or BNST oxytocin (p<0.01) neurons, aligning with observed measures in the literature (Chen et al. 2018; Popescu et al. 2017). The amplitude of sPSCs was also recorded across the same six populations, revealing significant differences in the size of individual synaptic events (**Fig. 4K**, F_5,102_=5.08, p=0.001). Notably, pPVN oxytocin neurons had significantly larger sPSC amplitudes compared to aPVN oxytocin neurons and BNST oxytocin neurons (p<0.01), suggesting that pPVN oxytocin neurons may receive stronger individual synaptic inputs.

We also assessed whole-cell membrane capacitance across populations. Cortical neurons had significantly higher Cm compared to aPVN oxytocin (**Fig. 4L**, p<0.0001) and BNST oxytocin neurons (p<0.0001). This is consistent with the larger somatodendritic size of cortical pyramidal neurons (Paramo et al. 2021). Resting membrane potential recordings also significantly varied across cell types (**Fig. 4M**, F_5,97_=9.71, p<0.001). Within the PVN, posterior oxytocin neurons were more hyperpolarized than anterior oxytocin neurons (p<0.001). Within the BNST, no significant differences were found between oxytocin neurons and non-oxytocin neurons. Cortical neurons were the most hyperpolarized (p<0.0001 vs. aPVN^OT^), aligning with related studies (Gniel & Martin 2010). Measurements of membrane resistance across cell types also showed differences (**Fig. 4N**, F_5,101_=9.75, p<0.0001). Posterior PVN oxytocin neurons had significantly lower membrane resistance than aPVN oxytocin neurons (p<0.0001), while PVN non-oxytocin neurons were intermediate, suggesting that aPVN oxytocin neurons have reduced leak conductance and may be more sensitive to synaptic input. In the BNST, membrane resistance was comparable between oxytocin and non-oxytocin neurons. Firing responses to stepwise current injection (0-100 pA) also revealed differences between the cell types (**Fig. 4O**, F_4,81_=22.49, p<0.001). Oxytocin neurons in the aPVN and BNST exhibited higher spike frequencies at all steps above 50 pA (cell type × current F_24,358_=8.88, p<0.001) compared to pPVN oxytocin neurons and non-oxytocin PVN neurons. Cortical neurons had the lowest overall excitability, replicating previous recordings (Eyal et al. 2018). Lastly, oxytocin neurons were held at -47 mV, closer to the reversal potential for chloride ions at physiological conditions (Lopantsev & Schwartzkroin 2001) and to the resting membrane potential for these cells (**Fig. 4M**). Both downward (sEPSCs) and upward (sIPSCs) were observed at this voltage in all cells (**Fig. 4P**). There were significantly more sEPSC events than sIPSC events in BNST (**Fig. 4Q**), aPVN (**Fig. 4R**), and pPVN (**Fig. 4S**) (all p<0.001), with pPVN receiving fewer sEPSCs and sIPSCs. These findings suggest regional differences of intrinsic excitability, with aPVN and BNST oxytocin neurons sharing a distinct, more excitable phenotype than pPVN oxytocin neurons.

## Discussion

There is growing evidence for heterogeneity in PVN oxytocin neurons, and we defined key anatomical distinctions within the PVN that help to explain how oxytocin produced in the PVN can drive opposing social behaviors. In California mice, we observed that oxytocin in the aPVN facilitates stress-induced social avoidance and vigilance, similar to BNST oxytocin neurons (Duque-Wilckens et al., 2020). In contrast, oxytocin produced in pPVN facilitates social approach in male and female California mice, similar to studies in other species showing that oxytocin promotes affiliative behaviors (Carcea et al., 2021; Dölen et al., 2013). We observed that the electrophysiological properties of aPVN oxytocin neurons in *Oxt*^Cre^ mice more closely resembled BNST oxytocin neurons as opposed to pPVN oxytocin neurons. Together, these results define a key functional heterogeneity within PVN oxytocin neurons and suggest that distinct populations of oxytocin neurons within the PVN enhance social salience in affiliative or aversive behavioral contexts.

### Oxytocin Produced in the Anterior PVN Promotes Avoidance and Vigilance

In female California mice, oxytocin neurons in the aPVN are more reactive during a social interaction test two weeks post-defeat compared to controls (Steinman et al., 2016). Our knockdown results suggest that this increased reactivity of aPVN oxytocin neurons contributes to stress induced social avoidance and vigilance in stressed female California mice. Others have also reported that aPVN oxytocin neurons do not promote affiliative responses. Both male and female wild house mice exhibit aggression in a resident-intruder test, and aggressive encounters increased oxytocin/c-fos colocalizations in aPVN oxytocin neurons in both sexes (Sofer et al., 2024). These neurons help to drive aggressive behavior in females. Ablation of oxytocin neurons decreased female aggression while optogenetic stimulation of these neurons increased aggression and arousal in females. Although oxytocin is usually considered to promote affiliation (Carter et al., 2008), it can also promote avoidance of unfamiliar individuals (Beery 2015). Thus, avoidance of unfamiliar individuals in pair bonded animals could be mediated by aPVN oxytocin neurons.

Although aPVN and pPVN differ in glutamatergic forebrain inputs (Ulrich-Lai et al. 2014), relatively little is known about what these oxytocin neuron populations may be encoding. It is likely that aPVN oxytocin neurons project to the anteromedial BNST because oxytocin receptor activation in this area promotes social avoidance and vigilance (Duque-Wilckens et al., 2018; Luo et al., 2022), and these two behaviors are reduced with oxytocin knockdown in the aPVN. Others have found a considerable density of PVN oxytocin fibers in the BNST (Knobloch et al. 2012). Future studies are needed to determine if these fibers originate primarily from the aPVN instead of pPVN.

### Oxytocin Produced in the Posterior PVN Promotes Social Approach

Our findings support the view that oxytocin neurons in the pPVN facilitate affiliative behaviors and motivation. While most studies do not explicitly target pPVN oxytocin neurons, several have focused on the roles of parvocellular oxytocin neurons. In *M. musculus* and Wistar rats, parvocellular neurons are concentrated in the pPVN (Lewis et al. 2020; Eliava et al. 2016). Although we did not determine whether our manipulations in California mice targeted magno-or parvocellular oxytocin neurons, our results are consistent with *Mus* studies showing that oxytocin produced in the pPVN promote affiliative social behaviors. This is interesting because there is strong evidence that parvocellular oxytocin neurons project to the mesolimbic dopamine system.

One study injected retrobeads into the mouse NAc and recorded from neurons that co-expressed retrobeads and oxytocin. The majority of recorded neurons exhibited parvocellular electrophysiological characteristics (Lewis, et al. 2020). This aligns with evidence linking oxytocin receptor activity in the NAc and VTA with social approach. For example, oxytocin antagonist in the NAc impairs social bonding in prairie voles (Liu and Wang, 2003) and social approach in California mice (Williams et al. 2020). Additionally, disrupting oxytocin signaling in the NAc abolished socially conditioned place preference (Dölen et al. 2013). The primary source of dopamine to the NAc is the VTA, and there is evidence that activation of oxytocin receptors in VTA has strong effects on social motivation as well (Shahrokh et al., 2010; Song et al., 2016). Oxytocin neurons in the pPVN also project to the VTA (Son et al. 2022). In *M. musculus*, while parvocellular oxytocin neurons are most prevalent in the pPVN, some magnocellular oxytocin neurons are intermixed with them. Tracing studies show that while most oxytocin neurons projecting to the VTA are parvocellular, a few are magnocellular (Hung et al. 2017). Thus, in addition to magnocellular oxytocin neurons showing similar morpho-electrical properties to parvocellular oxytocin neurons in the PVN (Chen et al., 2022), there may be some shared functional properties as well.

Notably, PVN oxytocin neurons send projections to both the NAc and VTA (Knobloch et al. 2012; Hung et al. 2017), and oxytocin receptors are expressed on dopaminergic neurons in the VTA that project to the NAc (Peris et al. 2017). This suggests a two-pronged circuit model of how oxytocin from the PVN modulates the VTA and NAc. Finally, we observed that oxytocin knockdown reduced social approach in stress naïve California mice but did not increase social vigilance. This result parallels findings of oxytocin receptor inhibition in the NAc, which reduces social approach but does not increase vigilance in stress-naïve California mice (Williams et al., 2020). These data support the hypothesis that social approach and vigilance are modulated by distinct but overlapping circuits.

### Electrophysiological Distinctions and Implications

The physiological differences between aPVN and pPVN oxytocin neurons support the idea that these populations are functionally distinct. Anterior PVN oxytocin neurons shared several key electrophysiological properties with oxytocin neurons in the BNST including significantly higher sPSC frequency and spike frequency. Both aPVN and BNST oxytocin neurons exhibited a longer latency to spike after current injections, resembling the A-type potassium current that is characteristic of magnocellular oxytocin neurons (Althammer & Grinevich, 2017). Posterior PVN oxytocin neurons had shorter latencies to spike, resembling the T-type potassium currents observed in parvocellular oxytocin neurons (Eliava et al. 2016). In *M. musculus*, there have been several reports that magnocellular oxytocin neurons dominate in the aPVN, while parvocellular are more prevalent in the pPVN (Lewis et al., 2020). Our electrophysiological results are consistent with these reports.

There were intriguing physiological similarities between aPVN and BNST oxytocin neurons. Our morpholino data indicate that aPVN oxytocin neurons, like BNST oxytocin neurons (Duque-Wilckens et al. 2020), drive stress-induced social avoidance and vigilance. An interesting possibility is that BNST and aPVN oxytocin neurons may share a cellular origin. Although the developmental origins of oxytocin neurons in the BNST are unknown, previous work indicates that BNST vasopressin neurons in rats are formed on embryonic day 12 (E12) and E13, while non-vasopressin cells originate between E14-E16 (Al-Shamma and De Vries, 1996). Similarly, the majority of magnocellular neurons (which includes oxytocin and vasopressin neurons) are generated between E12 and E13 in rats (Markakis and Swanson, 1997). This raises the possibility that oxytocin neurons outside of the hypothalamus, often referred to as the accessory nuclei (Althammer and Grinevich, 2018), could originate within the hypothalamus and migrate to their final location in the adult brain. There is precedent for this concept as gonadotropin releasing hormone neurons originate in the developing olfactory epithelium and migrate through the basal forebrain until reaching the hypothalamus (Tobet and Schwarting, 2006). It is important to note that BNST oxytocin neurons do not fully mimic the electrophysiological profile of the aPVN oxytocin neurons. This suggests that extrinsic factors such as cellular inputs or adjacent glial cells may impact the electrophysiological properties of BNST oxytocin neurons. Overall, our results showing similarities between aPVN and BNST oxytocin neurons suggest that future studies exploring the origin of extra-hypothalamic oxytocin neurons are warranted.

While we did not use tracing techniques to identify magnocellular oxytocin neurons, our observed physiological differences between aPVN and pPVN oxytocin neurons were largely consistent with previously identified characteristics of magno- and parvocellular oxytocin neurons (Sawchenko & Swanson, 1980; Eliava et al. 2016). However, there is growing evidence that a binary classification of oxytocin neurons may not fully capture observed complexity. Rigorous morpho-electric profiling revealed some overlap in physiological profiles in magno- and parvocellular neurons (Chen et al. 2022). An unsupervised clustering analysis of morpho-electric properties used action potential kinetics, dendritic complexity, soma size, and intrinsic excitability. This analysis created several clusters that included both magno- and parvocellular neurons. For example, magno- and parvocellular oxytocin neurons found in the pPVN were found to have similar characteristics. Additionally, along the anterior-posterior axis of the PVN, electrophysiological and morphological traits gradually shifted among magnocellular oxytocin neurons. Interestingly there is also evidence for coordination between magno- and parvocellular oxytocin neurons, as some parvocellular oxytocin neurons can modulate the activity of magnocellular oxytocin neurons in the PVN (Eliava et al. 2016). Together, these findings emphasize the concept that the form and function of oxytocin neurons changes across the anterior-posterior axis of the PVN.

## Conclusions

We observed that physiologically and functionally, aPVN oxytocin neurons had important differences from pPVN oxytocin neurons. In several dimensions, aPVN oxytocin neurons were more similar to BNST oxytocin neurons than pPVN oxytocin neurons, raising important questions about the cellular origin and development of these neurons. Previous tracing studies have focused primarily on differentiating between magnocellular and parvocellular neurons, with less attention paid to the relative location of these cells along the anterior-posterior axis. While recent morpho-electrical analyses suggested that this gradient is important for the physiological properties of PVN oxytocin neurons, it was unclear if this translated into functional differences. Our results suggest that indeed there are important functional differences between oxytocin produced in the anterior versus posterior PVN. Further study is needed to more fully characterize the projections of these neurons and to determine whether genetic markers can be identified that can predict the spatial location of oxytocin neurons within the hypothalamus.

## Acknowledgments

This work supported by T32 MH112507 to ANC, F31 MH134532 and T32 GM13432 to CS, NIMH R01 MH111604 to AJR, NIMH R01 MH121829 to BCT.

## Notes

### Competing Interest Statement

The authors have declared no competing interest.

### Summary of Updates

New photomicrographs, quantification of knockdown, and electrophysiological analyses added.

